# α-synuclein expression in response to bacterial ligands and metabolites in gut enteroendocrine cells

**DOI:** 10.1101/2023.04.02.535274

**Authors:** Michael J Hurley, Elisa Menozzi, Sofia Koletsi, Rachel Bates, Matthew E Gegg, Kai-Yin Chau, Hervé M Blottière, Jane Macnaughtan, Anthony H V Schapira

**Affiliations:** Department of Clinical and Movement Neurosciences, UCL Queen Square Institute of Neurology, London, UK; Institute for Liver and Digestive Health, University College London, London, UK; Université Paris-Saclay, INRAE, MetaGenoPolis, Jouy en Josas, & Nantes Université, INRAE, UMR 1280 PhAN, Nantes, France; Aligning Science Across Parkinson’s (ASAP) Collaborative Research Network, Chevy Chase, MD 20815, USA

**Keywords:** Toll-like receptor, free fatty acid receptor, short-chain fatty acid, enteroendocrine cells, microbiome, Parkinson disease

## Abstract

**Background:** Caudo-rostral migration of pathological forms of α-synuclein from the gut to the brain is proposed as an early feature in Parkinson disease (PD) pathogenesis, but the underlying mechanisms remain unknown. Intestinal enteroendocrine cells sense and respond to numerous luminal signals, including bacterial factors, and transmit this information to the brain via the enteric nervous system and vagus nerve. There is evidence that gut bacteria composition and their metabolites change in PD patients and these alterations can trigger α-synuclein pathology in animal models.

**Objective:** Here we investigated the effect of toll-like receptor (TLR) and free fatty acid receptor (FFA2/3) agonists on α-synuclein levels in mouse STC-1 enteroendocrine cells.

**Methods:** STC-1 cells were treated with TLR and FFA2/3 agonists alone and in combination with selective antagonists. The level of α-synuclein protein was measured in cell lysates and cell culture media by western blot and ELISA. And the level of α-synuclein and tumour necrosis factor (TNF) mRNA was measured by quantitative RT-PCR.

**Results:** TLR and FFA receptor agonists significantly increased intracellular and extracellular α-synuclein levels and antagonists significantly reduced these effects. TLR and FFA receptor agonists also significantly increased TNF transcription and this was inhibited by corresponding antagonists.

**Conclusions:** Elevated intracellular α-synuclein increases the likelihood of aggregation and conversion to toxic forms. Factors derived from bacteria induce α-synuclein accumulation in STC-1 cells. Here we provide support for a mechanism by which exposure of enteroendocrine cells to specific bacterial factors found in PD gut dysbiosis might facilitate accumulation and transmission of α-synuclein pathology from the gut to the brain.

## Introduction

Parkinson disease (PD) is a progressive neurodegenerative disorder clinically characterised by bradykinesia, rigidity, resting tremor with postural and gait abnormalities. Pathological features of PD include degeneration of midbrain dopaminergic neurons, brainstem catecholamine neurons and cholinergic neurons in the basal forebrain,^1,2^ together with deposits of aggregated α-synuclein in the central nervous system and periphery.^3-5^

While motor symptoms are the cardinal features of PD, non-motor symptoms adversely affect PD patients’ quality of life; among these, gastrointestinal disorders such as oropharyngeal/oesophageal dysphagia, gastroparesis and constipation frequently occur.^6^ ^7^ ^8^ The gastrointestinal tract may potentially play a central role in PD pathogenesis. For some individuals, spread of α-synuclein from the gastrointestinal tract to the brain could initiate the disease and underlie some of the manifestations affecting the gastrointestinal tract.^3^ ^4^ Indeed, different subtypes of PD (“gut-first” and “brain-first”) have been suggested. Within the former group, α-synuclein pathology might start in the periphery, within the enteric and parasympathetic nervous system, reflecting the presence of prodromal gastrointestinal disorders, and subsequently spread to the sympathetic nervous system and lower brainstem.^9^ To demonstrate a “gut-first” subtype, the trigger and location of the pathological cascade in the gastrointestinal tract in PD need to be elucidated. Mounting evidence suggests that the trillions of bacteria living in the gut, the gut microbiome, could initiate PD pathology. Experimentally, transfer of specific gut microbiome components and intestinal infection can cause Parkinson-like brain pathology in animal models^10-12^ and altered gut microbiome composition was found in PD patients^13-17^. Intestinal infection could therefore act as a catalyst to promote shifts in the microbiome composition, allowing gastrointestinal microbial products and inflammatory mediators to increase α-synuclein levels in the gut and subsequently the brain.^18^

A specific sub-population of intestinal epithelial cells – enteroendocrine cells – that are present in both small intestine and colon are considered a potential candidate for initial gut pathology in PD. Enteroendocrine cells comprise approximately 1 % of gut epithelium, express α-synuclein,^19^ and display neuronal-like properties including synaptic features^20^ and electrical excitability^21^. In contrast, α-synuclein is undetectable in enterocytes which make up the majority of the gut epithelium.^22^ Since enteroendocrine cells connect directly to enteric neurons and to the brainstem through the vagus nerve,^23^ they may act as a conduit for transfer of α-synuclein pathological forms from the gut lumen to brain via enteric innervation and vagus nerve.^19^ ^24^

Enteroendocrine cells express receptors for bacterial-derived components (toll-like receptors, TLR) and they respond to TLR agonists with increased proinflammatory cytokine production.^25-28^ Several lines of evidence have implicated a TLR-mediated response to bacterial ligands in PD pathogenesis. TLR4-knockout mice exhibit attenuated motor dysfunction and neuro-/gastrointestinal-inflammation and neurodegeneration in response to rotenone treatment.^29^ TLR2-deficient, α-synuclein over-expressing mice showed less neurodegeneration and neuronal α-synuclein concentrations.^30^

Enteroendocrine cells also express free fatty acids receptors (FFA) which recognise short-chain fatty acids (SCFA).^31^ ^32^ SCFA (mainly acetate, propionate and butyrate) are key bacterial metabolites promoting host intestinal health and they have been shown to modulate PD pathogenesis in different PD models with contrasting results.^11^ SCFA supplementation to germ-free α-synuclein-overexpressing mice reversed the neuroprotective effects of germ-free conditions suggesting a causal role,^11^ however sodium butyrate treatment ameliorated gut dysfunction and motor deficits in a rotenone-induced mouse model,^33^ and increased α-synuclein mRNA in STC-1 cells.^28^

Here we use mouse STC-1 enteroendocrine cells to assess the effect of treatment with TLR and FFA receptor agonists on α-synuclein expression and release.

## Materials and Methods

Unless otherwise stated chemicals were purchased from Merck and other reagents from Thermo Fisher Scientific.

### Mouse intestine

Ileal tissue from a 10 month *Gba* N370S colony wildtype mouse ^34^ was embedded in wax and sectioned. Animal husbandry and experimental procedures were performed in compliance with the UK Animal (Scientific Procedures) Act of 1986 at the UCL biological resources facility. <u>dx.doi.org/10.17504/protocols.io.eq2ly72ywlx9/v1</u>

### Cell culture

Mouse (*Mus musculus*) neuroendocrine duodenal adenoma STC-1 cells (RRID:CVCL_J405) were grown in DMEM:F12 Ham with GlutaMax^™^ medium supplemented with 10% charcoal absorbed foetal bovine serum (LabTech), 1 mM sodium pyruvate, non-essential amino acids (100 μM) and penicillin (100 U/ml) streptomycin (100 μg/ml) at 37°C under saturating humidity in a 5% CO_2_/95% air mixture. <u>dx.doi.org/10.17504/protocols.io.5jyl8jky8g2w/v1</u>

For immunohistochemistry cells were grown on 12 mm glass coverslips treated with poly-D-lysine or Geltrex^™^. After treatments, cells were washed with PBS, fixed with 4% paraformaldehyde for 15 min, washed twice with PBS and stored at -30°C. <u>dx.doi.org/10.17504/protocols.io.kqdg39yjpg25/v1</u>

Cell viability was assessed by Resazurin Reduction Assay (RRA) using a CellTiter-Blue^®^ Cell Viability Assay (Promega) according to the manufacturer’s instructions.

### Cell treatments

For TLR4 stimulation, cells were treated with 1 and 10 µg/ml of TLR4 selective ultrapure LPS-EB from *E. coli* 0111:B4 (LPS) (InvivoGen) for 24 or 48 hours. For TLR2 stimulation, cells were treated with 0.1 µg/ml and 1 µg/ml of Pam3CSK4 (PAM) (Cambridge Bioscience) for 24 or 48 hours. PAM is a selective ligand for the TLR2/TLR1 heterodimer.^35^ For antagonist experiments cells were pre-treated (1 hour) with TLR4 inhibitor TAK-242 (1 µM)^36^ before addition of 10 µg/ml LPS for 24 hours or the murine TLR2/1 signalling inhibitor C29 (50 µM)^37^ before addition of 1 µg/ml PAM for 24 hours. Control groups were treated with agonists or antagonists alone for 24 hours.

For FFA2/3 receptor stimulation, cells were treated with SCFA (4 mM sodium acetate (acetate), 4 mM sodium propionate (propionate), 2 mM sodium butyrate (butyrate)) for 24 hours. The effect of 1 hour pre-treatment with 1 µM GLPG0974 or 20 mM β-hydroxybutyrate on 24-hour butyrate stimulation was tested. GLPG0974 is a potent antagonist of the human FFA2 receptor but has lower affinity for rodent FFA2 isoforms ([^3^H]GLPG0974, K*_d_* 8.1 ± 0.9 and > 150 nM respectively).^38^ β-hydroxybutyrate is an FFA3 receptor antagonist.^39^ Control groups were treatment with antagonists or butyrate alone for 24 hours.

Following treatments cell culture medium was removed and frozen and cells were washed with phosphate buffered saline (PBS) and then frozen.

### Immunohistochemistry

Sections were processed for immunohistochemistry and staining visualised by immunofluorescence as previously described ^40^. Details of antibodies are given in **Table 1**. <u>dx.doi.org/10.17504/protocols.io.eq2ly72ywlx9/v1</u>

**Table 1.** Primers and Immunoglobulins

| Reagent | Catalogue # | Supplier | IHH/Western dilution | RRID<br>scicrunch.org/resources |
| --- | --- | --- | --- | --- |
| *Primary antibodies* |  |  |  |  |
| Rabbit anti-α-synuclein | ab212184 | www.abcam.com/ | 1:500/1:1250 | not available |
| Rabbit anti-β-actin | ab241153 | www.abcam.com/ | -/1:5000 | not available |
| Rat anti-5-HT | MAB352 | www.merckmillipore.com/ | 1:200/- | RRID:AB_94865 |
| Mouse anti-GLP-1 | ab26278 | www.abcam.com/ | 1:200/- | RRID:AB_470838 |
| Mouse anti-Cholecystokinin 8 | ab37274 | www.abcam.com/ | 1:200/- | RRID:AB_726010 |
| *Secondary antibodies* |  |  |  |  |
| Goat anti-rabbit IgG biotin | 111-065-144-JIR | https://strattech.com | 1:250/1:1000 | RRID:AB_2337965 |
| Goat anti-mouse IgG biotin | 115-065-003-JIR | https://strattech.com | 1:250/- | RRID:AB_2338557 |
| Goat anti-rat IgG biotin | 43R-1528-FIT | https://strattech.com | 1:250/- | RRID:AB_10814469 |
| Goat anti-mouse IgG Alexa Fluor™ Plus 555 | A32727 | www.thermofisher.com/ | 1:200/- | RRID:AB_2633276 |
| Goat anti-rabbit IgG Alexa Fluor™ 488 | A27034 | www.thermofisher.com/ | 1:200/- | RRID:AB_2536097 |
| Goat anti-rat IgG Alexa Fluor™ 546 | A-11081 | www.thermofisher.com/ | 1:200/- | RRID:AB_2534125 |
| Streptavidin POD conjugate | 11089153001 | sigmaaldrich.com | 1:250/1:1000 | not available |

IHH = immunohistochemistry; POD = peroxidase

Cells grown on glass coverslips were stained by the same procedure used for wax sections (excluding antigen retrieval) and visualised by 3,3′-Diaminobenzidine peroxidase reaction (**Table 1**). <u>dx.doi.org/10.17504/protocols.io.kqdg39yjpg25/v1</u>

Images were obtained using an Olympus BX43 microscope and AKOYA Biosciences Mantra II imaging system.

### Real-time quantitative PCR

Cells were processed to extract total RNA and protein using TRI Reagent^®^ solution and cDNA generated using a high-capacity cDNA reverse transcription kit and random primers. Quantitative real-time PCR was conducted with a StepOne Real-Time PCR system (Applied Biosystems) using PowerUp^™^ SYBR^™^ Green Master Mix and gene specific (BLASTN, RRID:SCR_001598) primers (**Table 1**) to quantify relative gene expression levels of α-synuclein, TNF and NF-κβ mRNA in the samples normalised to β-actin using the comparative 2^-ΔΔ^C_T_ method.^41^ <u>dx.doi.org/10.17504/protocols.io.rm7vzb2z2vx1/v1</u>

### Western blot

Protein pellets from the TRI Reagent^®^ extraction were dissolved in 2% SDS containing 8 M urea or cells were lysed in RIPA buffer containing protease inhibitors. Samples (∼ 10 ug total protein) were separated by polyacrylamide gel electrophoresis and protein transferred onto polyvinylidene difluoride membranes that were then probed with rabbit monoclonal antibodies against α-synuclein and β-actin overnight (**Table 1**). Bands were visualized by 3,3′-Diaminobenzidine/peroxidase reaction or with ECL plus reagents (Amersham), digitized using a ChemiDoc^™^ MP imaging system (Biorad) and analysed using ImageJ v1.53u (RRID:SCR_003070, https://imagej.net/) <u>dx.doi.org/10.17504/protocols.io.bp2l69pqdlqe/v1</u>

### Enzyme-linked immunosorbent assay

The amount of α-synuclein in the culture medium was measured using a mouse α-synuclein ELISA kit (Abcam ab282865) according to the manufacturer’s instructions.

### Proteosome assay

Ubiquitin-proteasome system (UPS) activation was assessed using a proteasome activity assay kit (Abcam ab107921) according to the manufacturer’s instructions.

### Data analysis

Data were analysed and visualized with GraphPad Prism^™^ v.9.5 (RRID:SCR_002798, http://www.graphpad.com/). Parametric data (Shapiro-Wilk, *P* > 0.05) were analysed by one-way ANOVA (*F*) and non-parametric data by Kruskal-Wallis one-way ANOVA by ranks (*H*). When ANOVA indicated a significant difference, *post hoc* multiple comparison pair-wise comparisons of all treatment groups or multiple comparisons of all treatment groups to the control group were made. Values are given as mean ± standard error of the mean. All tests were two-tailed and the significance level for all tests was taken to be *P* < 0.05.

## Results

### Enteroendocrine cells but not enterocytes express α-synuclein

Diffuse α-synuclein immunoreactivity and punctate cholecystokinin and glucagon-like peptide-1 (both released by and indicative of enteroendocrine cells) immunoreactivity was detected in STC-1 cells but staining for 5-hydroxytryptamine was not detectable (**Figure 1 A-D**).

**Figure 1.**
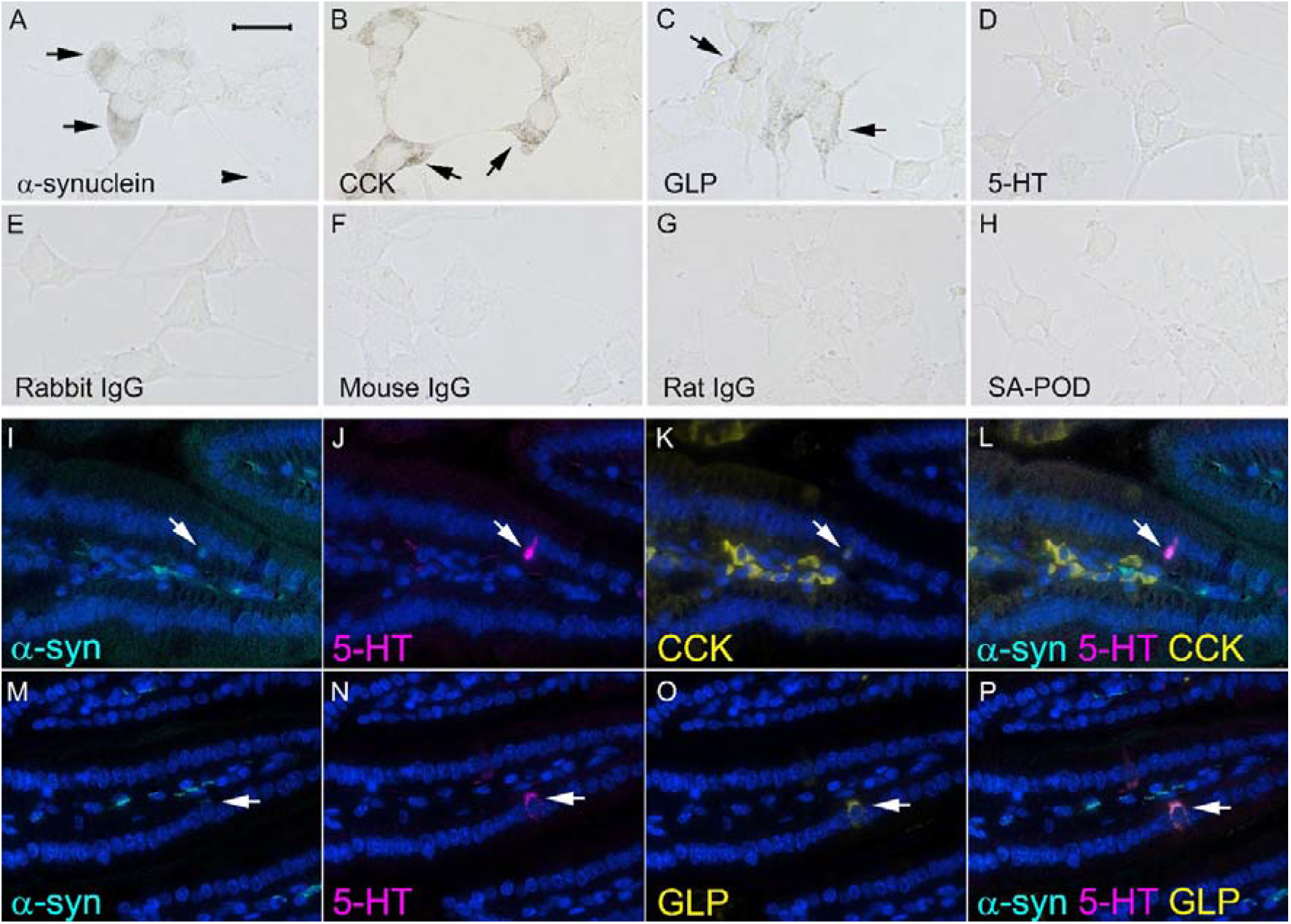
Co-expression of α-synuclein with enteroendocrine cell markers. Representative immunohistochemistry and immunofluorescence images showing immunoreactivity for α-synuclein and markers of enteroendocrine cells in STC-1 cells (A-H) and longitudinal sections of mouse ileum (I-P). (**A**) Diffuse α-synuclein immunoreactivity (arrows) was detected in STC-1 cells. The arrowhead shows a neuropod. (**B**) Punctate cholecystokinin (CCK) immunoreactivity (arrows) was detected in STC-1 cells, (**C**) Punctate glucagon-like peptide 1 (GLP) (arrows) was detected in STC-1 cells, (**D**) 5-hydroxytryptamine (5-HT) staining was not evident in STC-1 cells. No immunoreactivity was detected when primary antibodies were omitted and just biotinylated (**E**) rabbit (**F**) mouse or (**G**) rat immunoglobulins and streptavidin conjugated with peroxidase (SA-POD) were used, or (**H**) just SA-POD alone. (**I, M**) α-synuclein immunofluorescence (cyan) in the central neural tissue of a villus and an enteroendocrine cell (arrow) of mouse ileum. (**J, N**) 5-hydroxytryptamine (5-HT) immunofluorescence (magenta) of an enteroendocrine cell in mouse ileum, (**K**) cholecystokinin (CCK) immunofluorescence (yellow) in stromal cells of a villus and an enteroendocrine cell (arrow) of mouse ileum and (**O**) glucagon-like peptide (GLP) immunofluorescence (yellow) in an enteroendocrine cell (arrow) of mouse ileum. None of the markers (CCK, GLP, 5-HT) or α-synuclein were detectable in enterocytes. (**L**) co-expression of 5-HT, CCK and α-synuclein in an enteroendocrine cell of mouse ileum, (**P**) co-expression of 5-HT, GLP and α-synuclein in an enteroendocrine cell of mouse ileum. Cellular nuclei are stained blue with DAPI. Bar in panel A = 25 μm for panels A-H and 100 μm for panels I-P.

No staining was observed when primary antibodies were omitted and just biotinylated secondary antibody and streptavidin-peroxidase were used for the immunohistochemistry procedure (**Figure 1 E-G**). No staining was seen when just streptavidin alone was incubated with the cells (**Figure 1 H**).

Enteroendocrine cells within the epithelium of mouse ileum expressed α-synuclein, cholecystokinin, glucagon-like peptide-1 and 5-hydroxytryptamine, whereas α-synuclein was not detectable in ileal enterocytes (**Figure 1 I-P**).

### Effect of TLR and FFA2/3 agonists on α-synuclein

STC-1 cells constitutively express and release α-synuclein and this was augmented by TLR and FFA2/3 agonists (**Figure 2, 3**). Treatment with TLR agonists had no effect on cell viability (**Supplementary figure 1**).

**Figure 2.**
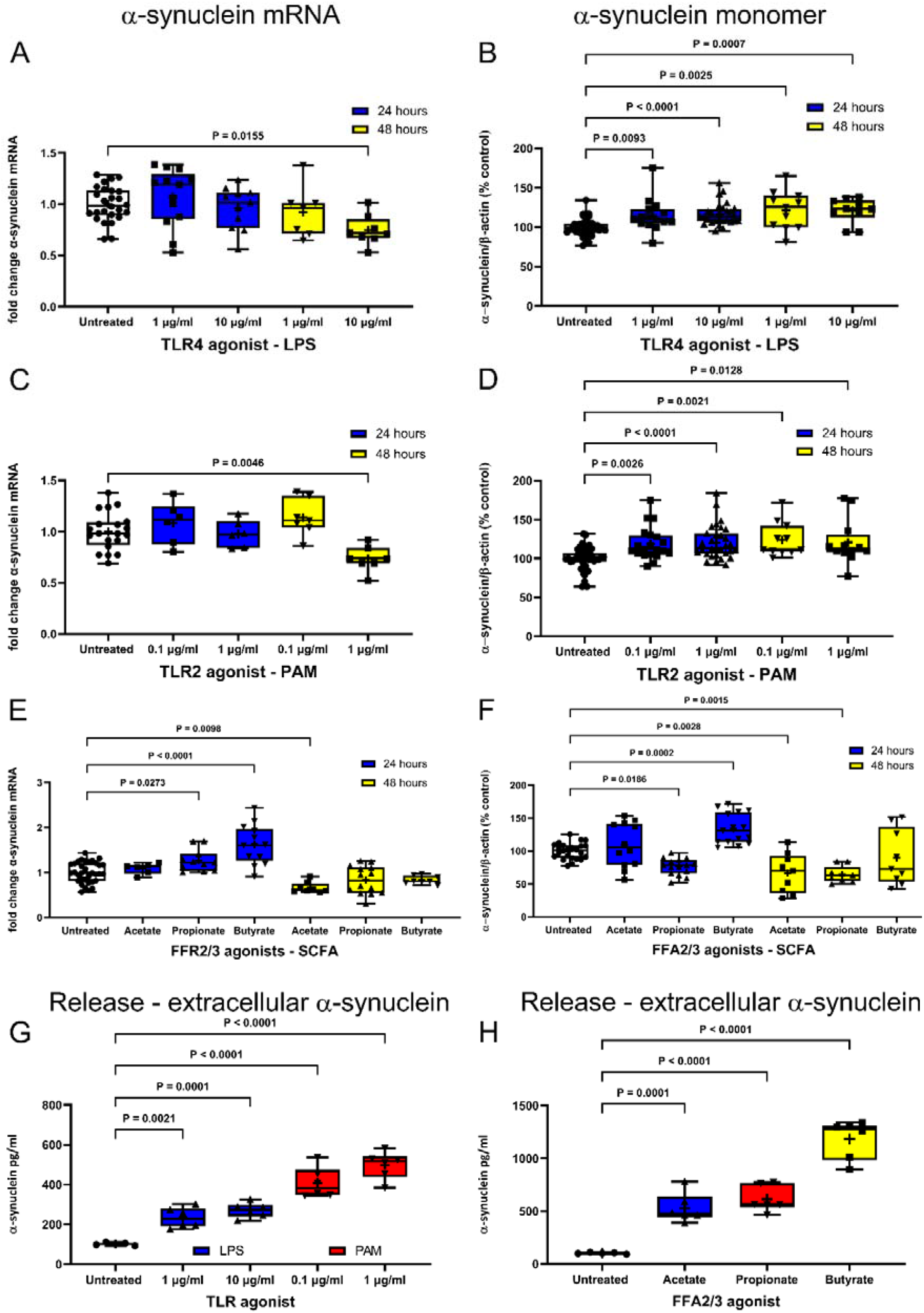
TLR and FFA2/3 agonists alter α-synuclein expression and release. LPS and PAM treatment of STC-1 cells decreased the level of α-synuclein mRNA after 48 hours but increased intracellular levels of α-synuclein monomer. Short chain fatty acids had opposing effects on α-synuclein mRNA and monomer. (**A**) 10 μg/ml LPS caused a 25 % reduction (*P* = 0.0155) in the level of α-synuclein mRNA compared to untreated cells after 48 hours. (**B**) 24-hour treatment with 1 and 10 μg/ml LPS caused a significant 15% (*P* = 0.0093) and 17% (*P* < 0.0001) increase in α-synuclein monomer respectively. 48-hour treatment with 1 and 10 μg/ml LPS caused a significant 22% (*P* = 0.0025) and 18 % (*P* = 0.0007) increase in α-synuclein monomer respectively. (**C**) 1 μg/ml PAM caused a significant 26 % reduction (*P* = 0.0046) in the level of α-synuclein mRNA compared to untreated cells after 48 hours. (**D**) 24-hour treatment with 0.1 and 1 μg/ml PAM caused a significant 16 % (*P* = 0.0026) and 19 % (*P* < 0.0001) increase in α-synuclein monomer respectively. 48-hour treatment with 0.1 and 1 μg/ml PAM caused a significant 40 % (*P* = 0.0021) and 28 % (*P* = 0.0128) increase in α-synuclein monomer. (**E**) 2 mM butyrate and 4 mM propionate increased α-synuclein mRNA after 24 hours by 57 % (*P* < 0.0001) and 26 % (*P* = 0.0273) and 4 mM acetate reduction α-synuclein mRNA by 36% (*P* = 0.0098) compared to untreated cells after 48 hours. (**F**) After 24 hours 2 mM butyrate increased α-synuclein monomer by 35 % (*P* < 0.0002) whereas 4 mM propionate decreased α-synuclein monomer by 31 % (*P* < 0.0186). After 48 hours 4 mM acetate and 4 mM propionate caused a 34 % (*P* = 0.0028) and 32 % (*P* = 0.0015) reduction in α-synuclein monomer respectively. TLR and FFA2/3 agonists increased release of α-synuclein protein from STC-1 cells. (**G**) Stimulation of STC-1 cells with 1 μg/ml and 10 μg/ml LPS for 24 hours increased α-synuclein protein release by 2.3 ± 0.2-fold (*P* = 0.0021) and 2.7 ± 0.2-fold (*P* < 0.0001) respectively and stimulation of STC-1 cells with 0.1 μg/ml and 1 μg/ml PAM for 24 hours increased α-synuclein protein release by 4.0 ± 0.3 -fold (*P* = 0.0001) and 4.9 ± 0.3-fold (*P* < 0.0001) respectively. (**H**) Stimulation of STC-1 cells with 4 mM acetate, 4 mM propionate or 2 mM butyrate for 24 hours increased α-synuclein protein release by 4.6 ± 0.5-fold (*P* = 0.0001), 5.4 ± 0.5 -fold (*P* < 0.0001) and 10.3 ± 0.7 -fold (*P* = 0.0001) respectively. Data are presented as box (25^th^ and 75^th^ percentiles) and whisker (5^th^ and 95^th^ percentiles), median (line), mean (+) with all data points. For qPCR data represent fold change levels of α-synuclein mRNA compared to untreated cells normalised to β-actin using the 2^-ΔΔ^C_T_ method. For western blot data represent relative levels of α-synuclein protein in comparison to untreated cells normalised to β-actin. For ELISA data represent fold change levels of α-synuclein compared to untreated cells. Data were analysed with Dunnett’s or Dunn’s multiple comparison test.

**Figure 3.**
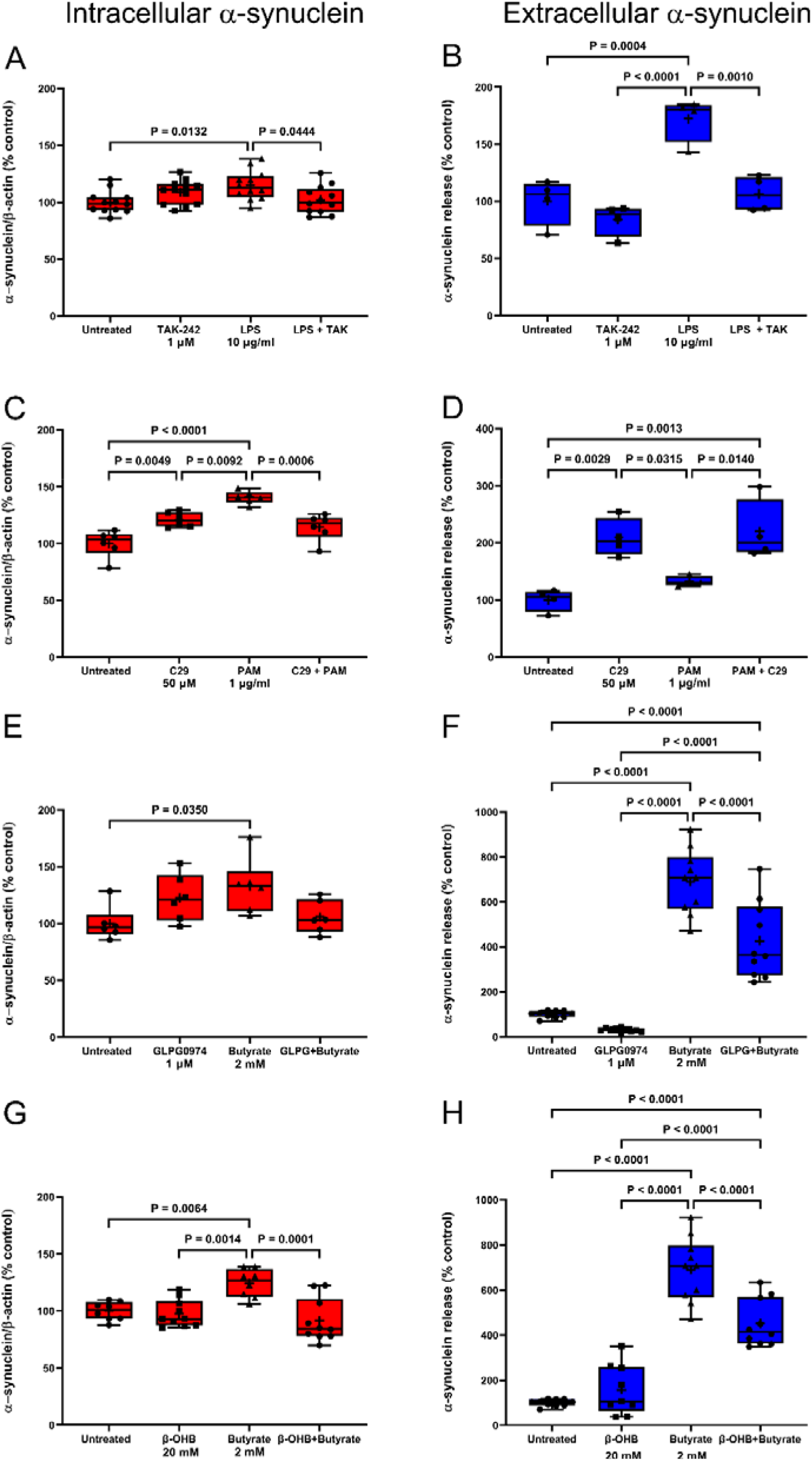
TLR and FFA2/3 antagonists reduce agonist-induced α-synuclein accumulation and release. Antagonists selective for TLR4, TLR2, FFA2 and FFA3 receptors significantly reduced the effect of their respective agonists on intracellular α-synuclein monomer content and α-synuclein release. Stimulation of STC-1 cells with 10 μg/ml LPS for 24 hours (**A**) increased intracellular α-synuclein monomer content by 15 % (*P* = 0.0132) which was significantly reduced (*P* = 0.0444) by 1 μM TAK-242 and (**B**) increased α-synuclein release by 75 % (*P* = 0.0004) and this was significantly reduced (*P* = 0.0010) by 1 μM TAK-242. TAK-242 treatment alone did no alter the intracellular α-synuclein monomer level or release of α-synuclein. Stimulation of STC-1 cells with 1 μg/ml PAM for 24 hours (**C**) increased intracellular α-synuclein monomer content by 41 % (*P* = 0.0001) and this was significantly reduced (*P* = 0.0006) by 50 μM C29. Treatment with C29 alone also caused a 21 % increase (*P* = 0.0049) in intracellular α-synuclein monomer. (**D**) Stimulation of STC-1 cells with 1 μg/ml PAM for 24 hours did not increase α-synuclein release, but C29 alone did cause a significant 109 % (*P* = 0.0029) increase in α-synuclein release. Stimulation of STC-1 cells with 2 mM butyrate for 24 hours (**E**) increased α-synuclein monomer content by 33 % (*P* = 0.0350) in comparison to untreated cells and this effect was mitigated by 1 μM GLPG0974 and (**F**) increased α-synuclein release by 690 % (*P* = 0.0001) and this was significantly reduced (*P* = 0.0001) by 1 μM GLPG0974. GLPG0974 alone did not alter the intracellular α-synuclein monomer level or release of α-synuclein. Stimulation of STC-1 cells with 2 mM butyrate for 24 hours (**G**) increased α-synuclein monomer content by 24 % (*P* = 0.0064) and this was significantly reduced (*P* = 0.0001) by 20 mM β-hydroxybutyrate and (**H**) increased α-synuclein release by 690 % (*P* = 0.0001) and this was significantly reduced (*P* = 0.0001) by 20 mM β-hydroxybutyrate. β-hydroxybutyrate alone had no significant effect on intracellular α-synuclein monomer content or α-synuclein release. Data are presented as box (25^th^ and 75^th^ percentiles) and whisker (5^th^ and 95^th^ percentiles), median (line), mean (+) and all data points and was analysed by one-way ANOVA and *post-hoc* Tukey’s multiple comparison test.

A representative western blot is shown in **Supplementary Figure 2**. Data for each graph and images of western blots can be found at the zenodo data repository.

(https://doi.org/10.5281/zenodo.7712730)

### α-synuclein expression

LPS 1 μg/ml and 10 μg/ml had no effect on the level of α-synuclein mRNA in comparison to untreated cells after 6 hours (data not shown) or 24 hours, but after 48 hours the higher concentration of LPS caused a significant reduction in the α-synuclein mRNA level (*F* (4, 57) = 4.168, *P* = 0.0050) (**Figure 2 A**). LPS 1 μg/ml and 10 μg/ml significantly increased total intracellular α-synuclein monomer levels after 24- and 48-hours (*H* (4, 101) = 14.29, *P* = 0.0001) treatment (**Figure 2 D**).

PAM 0.1 μg/ml and 1 μg/ml had no effect after 6 hours (data not shown) or 24 hours, but after 48 hours the higher concentration caused a significant reduction in the α-synuclein mRNA level (*F* (4, 40) = 6.143, *P* = 0.0006) (**Figure 2 B**). PAM 0.1 μg/ml and 1 μg/ml significantly increased total intracellular α-synuclein monomer levels (*F* (4, 99) = 9.621, *P* = 0.0001) after 24- and 48-hours treatment (**Figure 2 E**).

Stimulation of cells with butyrate and propionate for 24 hours significantly increased the level of α-synuclein mRNA, while acetate had no effect. In contrast, after 48 hours butyrate and propionate had no effect on α-synuclein mRNA whereas acetate caused a significant reduction (*F* (6, 90) 22.04, *P* = 0.0001) (**Figure 2 C**). Stimulation of cells with butyrate and propionate for 24 hours significantly increased and decreased respectively the level of intracellular α-synuclein monomer, while acetate had no effect. In contrast, after 48 hours butyrate had no effect on the level of α-synuclein monomer whereas acetate and propionate caused a significant reduction (*F* (6, 125) 23.03, *P* = 0.0001) (**Figure 2 F**).

### α-synuclein release

Untreated STC-1 cells released α-synuclein into culture medium that resulted in a concentration of 0.11 ± .01 ng/ml in a 24-hour period. LPS 1 μg/ml and 10 μg/ml for 24 hours caused a significant (*H* (2, 15) = 12.12, *P* < 0.0002) increase in the amount of released α-synuclein to 0.23 ± 0.05 ng/ml and 0.27 ± 0.02 ng/ml respectively (**Figure 2 G**). PAM 0.1 μg/ml and 1 μg/ml for 24 hours caused a significant (*H* (2, 15) = 12.78, P < 0.0001) increase in the amount of released α-synuclein to 0.41 ± .03 ng/ml and 0.50 ± 0.03 ng/ml respectively (Figure 2 G).

Stimulation of cells with acetate, propionate or butyrate for 24 hours caused a significant (*F* (3, 19) = 60.44, *P* < 0.0001) increase in the amount of released α-synuclein to 0.53 ± 0.06 ng/ml, 0.61 ± 0.05 ng/ml and 1.18 ± 0.08 ng/ml respectively (**Figure 2 H**).

### TLR antagonists reduce agonist-induced α-synuclein accumulation and release

Selective antagonists were used to demonstrate that agonist effects were receptor-mediated and not due to non-specific actions. The higher concentration of each TLR agonist was chosen because these gave the greatest and most robust response. TAK-242 reduced the increase in α-synuclein monomer (*F* (3, 44) = 4.214, *P* = 0.0105) and release (*F* (3, 12) = 19.54, *P* < 0.0001) induced by LPS after 24 hours, with no significant effect of TAK-242 treatment alone (**Figure 3 A, B**).

Similarly, C29 reduced the increase in α-synuclein monomer (*F* (3, 20) = 19.13, *P* < 0.0001) induced by PAM after 24 hours but had no effect on release. However, C29 treatment alone increased intracellular α-synuclein monomer (*F* (3, 20) = 19.13, *P* < 0.0001) and release (*F* (3, 12) = 12.24, *P* = 0.0006). PAM did not cause significant α-synuclein release in this experiment when analysed with Tukey’s multiple comparison test, however the difference between untreated and PAM treated cells was significant when analysed by an unpaired two-tailed t-test (t = 3.087, df = 6, *P* = 0.0215) (**Figure 3 C, D**).

### FFA2/3 antagonists reduce butyrate-induced α-synuclein accumulation and release

GLPG0974 reduced the increase in α-synuclein monomer (*F* (3, 20) = 3.744, *P* = 0.0277) and release (*F* (3, 36) = 75.48, *P* < 0.0001) induced by butyrate. GLPG0974 treatment alone had no significant effect on intracellular α-synuclein levels or release (**Figure 3 E, F**).

β-hydroxybutyrate significantly reduced the increase in α-synuclein monomer (*F* (3, 32) = 9.396, *P* < 0.0001) and release (*F* (3, 35) = 68.05, *P* < 0.0001) induced by butyrate, whereas β-hydroxybutyrate treatment alone had no significant effect on the level or release of α-synuclein (**Figure 3 G, H**).

### TLR and FFA2/3 agonists stimulate TNF mRNA production

The inflammatory consequences of TLR and FFA2/3 activation were investigated. Although levels of NF-κβ mRNA were not influenced by any agonists at the 24-hour timepoint examined (**Supplementary Figure 3**), a significant increase in the level of TNF mRNA was observed after stimulation with 10 μg/ml LPS and 1 μg/ml PAM (*H* (4, 65) = 40.54, *P* < 0.0001) (**Figure 4 A**). Likewise, propionate and butyrate significantly increased TNF mRNA (*F* (3, 23) = 55.69, *P* < 0.0001) whereas acetate had no effect (**Figure 4 B**).

**Figure 4.**
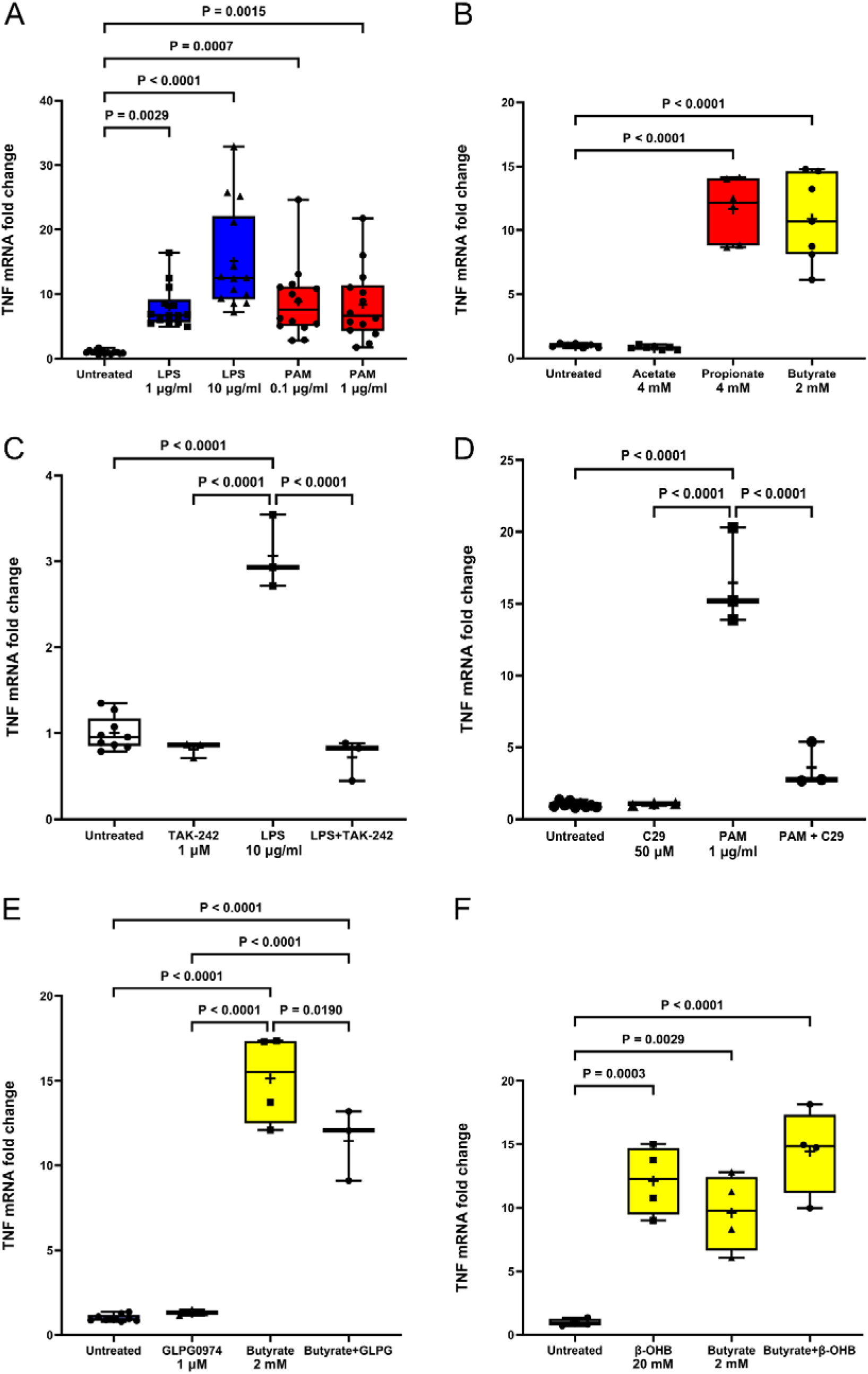
TLR and FFA2/3 agonists increase TNF mRNA in STC-1 cells. LPS, PAM and SCFA treatment of STC-1 cells significantly increased the amount of TNF mRNA. (**A**) Stimulation of STC-1 cells with 1 μg/ml and 10 μg/ml LPS for 24 hours increased TNF mRNA by 6.9-fold (*P* = 0.0029) and 16.2-fold (*P* < 0.0001) respectively and stimulation of cells with 0.1 μg/ml and 1 μg/ml PAM for 24 hours increased TNF mRNA by 7.9-fold (*P* = 0.0007) and 7.8-fold (*P* = 0.0015) respectively. (**B**) Stimulation of STC-1 cells with 4 mM acetate had no effect on TNF mRNA, whereas 4 mM propionate and 2 mM butyrate increased TNF mRNA by 11.7-fold (*P* < 0.0001) and 12.2-fold (*P* < 0.0001) respectively. Pre-treatment (1 hour) with: (**C**) TAK-242 (1 µM) significantly reduced (*P* < 0.0001) the LPS-induced increase in TNF mRNA, (**D**) C29 (50 µM) significantly reduced (*P* < 0.0001) the PAM-induced increase in TNF mRNA, (**E**) GLPG0974 (1 µM) significantly reduced (*P* < 0.0190) the butyrate-induced increase in TNF mRNA and (**F**) β-hydroxybutyrate (β-OHB) (20 mM) had no effect on the butyrate-induced increase in TNF mRNA but did itself cause a significant (*P* = 0.0003) increase in TNF mRNA. Data are presented as box (25^th^ and 75^th^ percentiles) and whisker (5^th^ and 95^th^ percentiles), median (line), mean (+) with all data points and represent relative levels of TNF-α mRNA compared to untreated cells and normalised to β-actin using the 2^-ΔΔ^C_T_ method. Data were analysed with Dunnett’s, Tukey’s or Dunn’s multiple comparison test.

The induction of TNF following LPS, PAM and butyrate exposure for 24 hours was significantly reduced by TAK-242 (*F* (3, 14) 68.72, *P* < 0.0001), C29 (*H* (3, 14) = 12.16, *P* = 0.0003) and GLPG0974 (*F* (3, 15) = 117.6, *P* < 0.0001) respectively, confirming specific activation of TLR4, TLR2 and FFA2 and involvement in stimulation of TNF transcription (**Figure 4 C-E**). β-hydroxybutyrate did not affect butyrate-induced TNF mRNA transcription, suggesting that the action of butyrate on TNF mRNA transcription was not through FFA3 receptors. However, β-hydroxybutyrate alone caused a significant increase in TNF mRNA (*F* (3, 12) = 19.74, P < 0.0001) (**Figure 4 F**).

### Ubiquitin-proteosome system

The UPS pathway was evaluated in cells stimulated with LPS 1 μg/ml, or PAM 0.1 μg/ml for 24 and 48h. Compared to untreated cells, there was a significant increase in proteasome activity in cells stimulated with either LPS or PAM at both time points (*F* (4, 13) = 21.93, *P* < 0.001) (**Supplementary Figure 4**), indicating that UPS impairment was not responsible for α-synuclein accumulation upon TLR activation.

## Discussion

The hypothesis that structural bacterial components and/or bacterial metabolites, such as SCFA, could alter α-synuclein expression in enteroendocrine cells was tested. Stimulation of enteroendocrine cells with TLR2 and TLR4 agonists, a lipoprotein and LPS respectively, increased intracellular α-synuclein monomer levels and release of α-synuclein into the culture medium, and that this effect was reduced by TLR antagonists. Activation of enteroendocrine cells with SCFA, particularly butyrate, increased release of α-synuclein into the culture medium. Again, the effect of butyrate was reduced by FFA2 and FFA3 receptor antagonists. Interestingly, butyrate and propionate had an opposing effect on intracellular α-synuclein levels. This may be due to the different affinities of each SCFA for the FFA2 and 3 receptors, the additional action of butyrate as a histone deacetylase inhibitor or other non-FFA receptor actions.^42^ Selective FFA receptor agonists or the histone deacetylase inhibitor trichostatin A could be used to investigate this further. Overall, these results suggest that enteroendocrine cells could be the initial site of α-synuclein accumulation in PD when stimulated by specific bacterial ligands through TLR2/4 activation and SCFA acting on FFA2/3.

It has been proposed that the pathological changes in a “body-first” PD subtype might begin within the gastrointestinal tract. The factors that can initiate these changes and where in the gastrointestinal tract they might occur, are unknown.^9^ Exposure to bacterial components, such as LPS for Gram-negative and a synthetic lipoprotein PAM for Gram-positive bacteria, increased intracellular accumulation and release of α-synuclein. The observed changes seemed to be specific to TLR4 and TLR2 activation as shown by the effective inhibition of LPS- and PAM-induced α-synuclein protein increase upon treatment with their respective antagonists. Activation of TLR4 with LPS and TLR2 with PAM resulted in the production of mRNA for the inflammatory cytokine TNF. Again, this effect on TNF was significantly reduced using selective antagonists. Butyrate also caused a large increase in TNF mRNA and this effect was not completely abolished by GLPG0974 (or β-hydroxybutyrate) suggestive of an additional mechanism to FFA activation for the induction of TNF mRNA by butyrate. The level of mRNA for the inflammatory transcription factor NF-κβ was not altered at the 24-hour timepoint tested which may reflect the more immediate response of transcription factors to inflammatory challenges and earlier timepoints or the nuclear trans-localisation of NF-κβ could be examined to see if NF-κβ is affected by TLR and FFA agonists.

Bacterial lipoproteins present in Gram-positive bacteria are recognised by TLR2 of the innate immune system that play a central role in the immune response to pathogenicity of bacterial species^43^ and TLR2 may have a role in PD pathogenesis. TLR2 protein was increased in post-mortem brain from patients with PD and positively correlated with disease status and duration.^44^ Knocking out TLR2 in transgenic mice expressing human A53T α-synuclein reduced levels of α-synuclein.^30^ *In vitro*, differentiated SH-SY5Y transduced with lentiviral α-synuclein and treated with PAM showed increased levels of insoluble α-synuclein oligomers. ^30^ Moreover, oligomeric forms of neuron-released α-synuclein activated microglial TLR2.^45^ Interestingly, small molecules inhibiting the TLR signal pathway ameliorated the PAM-induced increase of both TLR2 and α-synuclein in differentiated SHSY5Y cells, suggesting a possible therapeutic application of TLR2-targeting agents in PD.^44^ For the first time, we have analysed the role of TLR2 pathway in enteroendocrine cells and provided evidence supporting the link between TLR2 and PD pathogenesis in response to bacterial components. Application of TLR2 inhibitors or antagonists is of interest to mitigate pathogenic changes in the gut. The impact of TLR2 inhibitors should, however, be particularly considered in relation to those Gram-positive bacteria within the phylum *Firmicutes* which are particularly rich in lipoproteins. Among those, there is *Enterococcus faecalis*, an opportunistic pathogen responsible for nosocomial infections,^46^ and *Enterococcus faecalis* is one of the key bacteria affecting levodopa bioavailability.^47^ For these reasons, within PD, suppressing host TLR activity should be carefully considered not only for the well-known increased vulnerability to infection,^48^ but also for the potential impact on certain bacterial niches and its implication on disease course.

Neuroinflammation is implicated in PD pathogenesis and several models have used LPS to mimic the inflammatory events seen in PD.^49^ PD patients have increased expression of TLR4, enhanced markers of bacterial translocation and higher pro-inflammatory gene profiles in the colon when compared with healthy controls.^29^ In TLR4 knock-out mice, rotenone induced less intestinal inflammation, neuroinflammation and motor dysfunction compared to wild-type animals.^29^ Could treatments modulating the TLR4 inflammatory response ameliorate PD? In our study TAK-242, which is already under investigation in liver disease (<u>NCT04620148</u>), reduced the LPS-induced accumulation of α-synuclein in the enteroendocrine cells and testing such drugs in LPS models of PD is warranted.

Recent studies evaluating the gut microbiome in PD patients found an increase of opportunistic pathogens (*Porphyromonas*, *Prevotella*, *Corynebacterium 1*).^50^ Hence, we can hypothesise that these commensal bacteria could become prevalent in certain individuals and thus induce pathological changes seen in PD.

The same study also found a reduction of SCFA-producing bacteria in the PD population,^50^ which has been confirmed in other patient cohorts with reduced levels of SCFA detected in stool samples.^51^ This suggests that loss of SCFA in the lower GI tract might contribute to pathological manifestations and symptoms in PD. Our results showing increased α-synuclein expression and release in STC-1 induced by SCFA seem in contrast with these findings. However, STC-1 cells are derived from the duodenum and, to our knowledge, there is no study which has evaluated the composition of the microbiome in the small intestine. Duodenal cells would not normally be exposed to high levels of SCFA since they are produced in the colon in healthy individuals. However, people with PD suffer from small intestinal bacterial overgrowth^52^ and which bacterial species predominant in the upper GI tract of PD patients and their effect on enteroendocrine cells is unknown.

There are some limitations associated with this work. These experiments do not address the pathogenic mechanisms underlying α-synuclein accumulation and/or release (e.g., free or in exosomes) or where in the cell it accumulates (cytosolic or vesicular). Moreover, only the monomeric form of α-synuclein was analysed. Nevertheless, we show for the first time that LPS, lipoproteins and SCFA trigger α-synuclein pathology in enteroendocrine cells. Antagonists to the TLR and FFA receptors reversed the pathological changes and might become targets of interest for research into future treatments. Also, and of particular importance in relation to spread of α-synuclein pathology from gut to brain in PD is where from the cell α-synuclein is released. Enteroendocrine cells are polar epithelial cells and determining whether the release of α-synuclein is global, into the gut lumen from the apical surface or through synaptic connections onto cells of enteric nervous system is important to establish.

## Abbreviations

FFA: free fatty acid receptor
GLPG0974: 4-[[1-(benzo[b]thiophene-3-carbonyl)-2-methyl-azetidine-2-carbonyl]-(3-chloro-benzyl)-amino]-butyric acid
LPS: lipopolysaccharide
PAM: Pam3CysSerLys4
PBS: phosphate buffered saline
PBST: PBS containing 0.01% (v/v) Tween™-20)
PD: Parkinson disease
SCFA: short chain fatty acid
STC-1: mouse enteroendocrine cell line
TLR: toll-like receptor
TNF: tumour necrosis factor
UPS: ubiquitin-proteasome system

## Data availability

Data and methods that support the findings of this study are openly available at https://zenodo.org and https://www.protocols.io repositories respectively as referenced via DOI numbers throughout this manuscript.

## Author roles

(1) Research Project: A. Acquisition of Funding, B. Conceptualisation, C. Execution; (2) Statistical Analysis: A. Design, B. Execution, C. Review and Critique; (3) Manuscript: A. Writing of the First Draft, B. Review and Critique. Michael Hurley 1B, 1C, 2A, 2B, 2C, 3A, 3B; Elisa Menozzi: 1B, 1C, 2A, 2B, 2C, 3A, 3B; Sofia Koletsi: 1C, 3B; Rachel Bates: 1C, 3B; Matt Gegg: 1B, 3B; Kai-Yin Chau: 1B, 3B; Hervé Blottière 3B; Jane Macnaughtan: 1A, 1B, 3B; Anthony Schapira: 1A, 1B, 3B

## Financial disclosures of all authors (for the preceding 12 months)

The authors have no financial disclosures to declare for the preceding 12 months.

## Acknowledgements

This research was funded by Aligning Science Across Parkinson’s [Grant number: ASAP-000420] through the Michael J. Fox Foundation for Parkinson’s Research (MJFF). Elisa Menozzi is supported by a Royal Free Charity Fellowship. For the purpose of open access, the author has applied a CC BY 4.0 public copyright license to all Author Accepted Manuscripts arising from this submission.

---

**Supplementary Figure 1.**
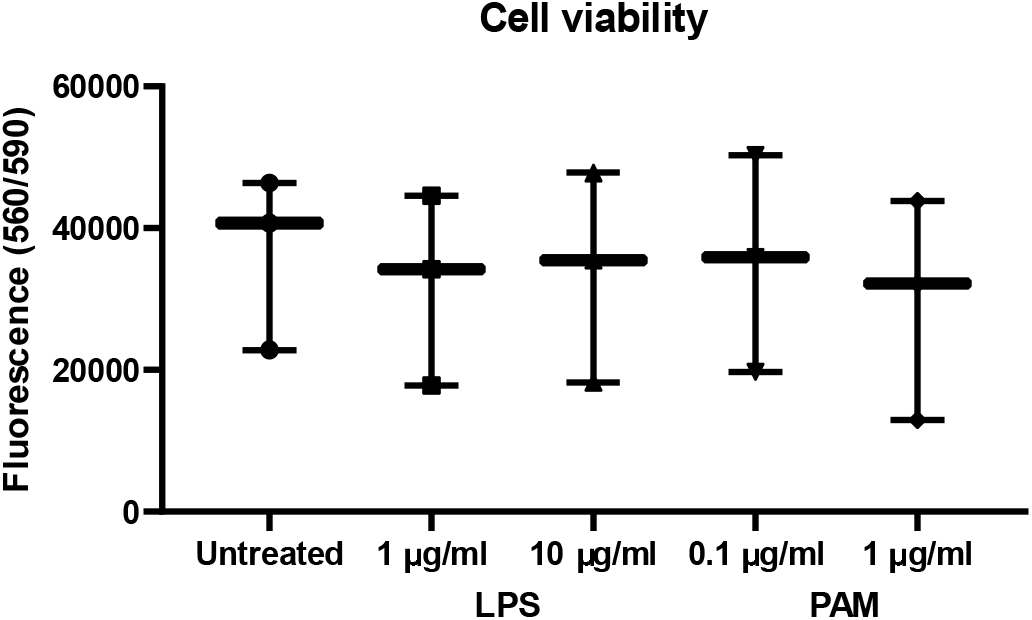
No differences in the number of viable cells were found after 24 hours exposure of STC-1 cells to two concentrations of LPS and PAM (Dunnett’s multiple comparison). Data are presented as box (25^th^ and 75^th^ percentiles) and whisker (5^th^ and 95^th^ percentiles), median (line), mean (+) and all data points.

**Supplementary Figure 2.**
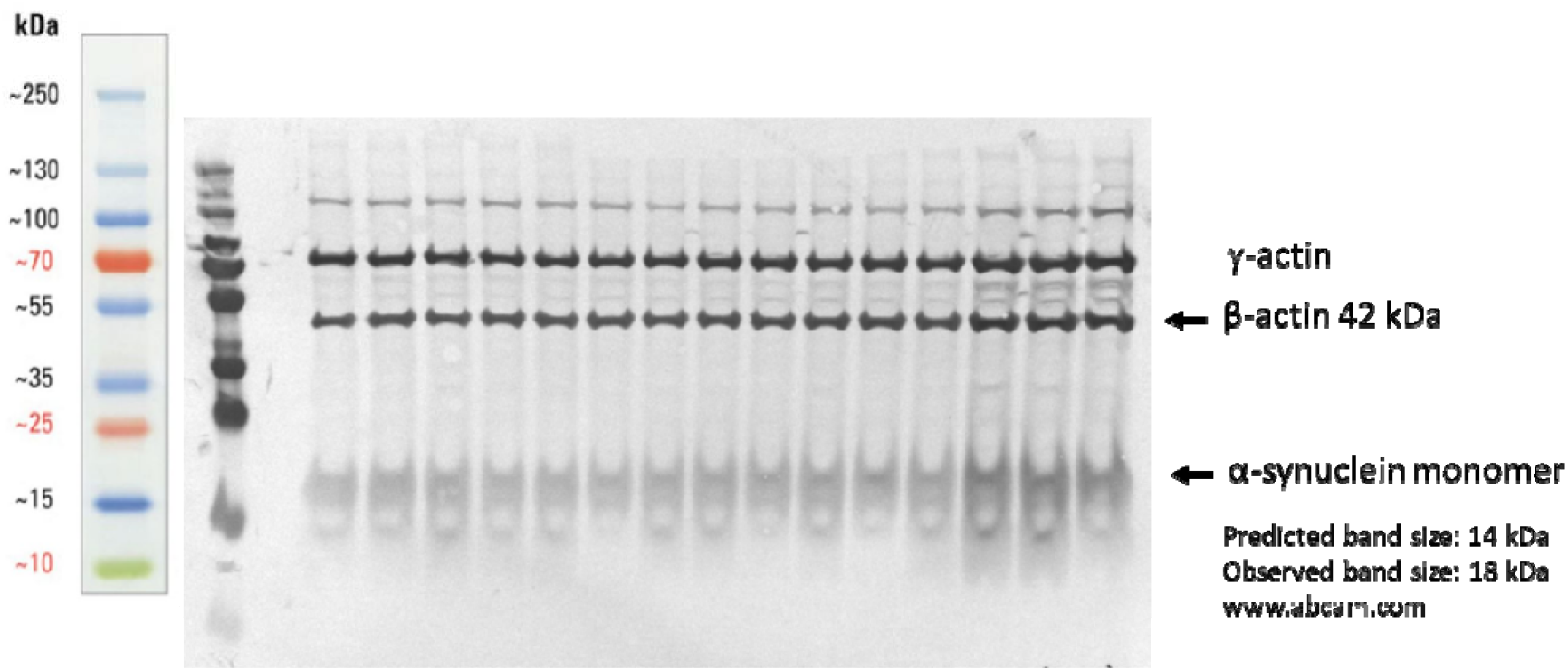
Representative blot used for semi-quantitative measurement of the level of α-synuclein protein monomer in STC-1 cells.

**Supplementary Figure 3.**
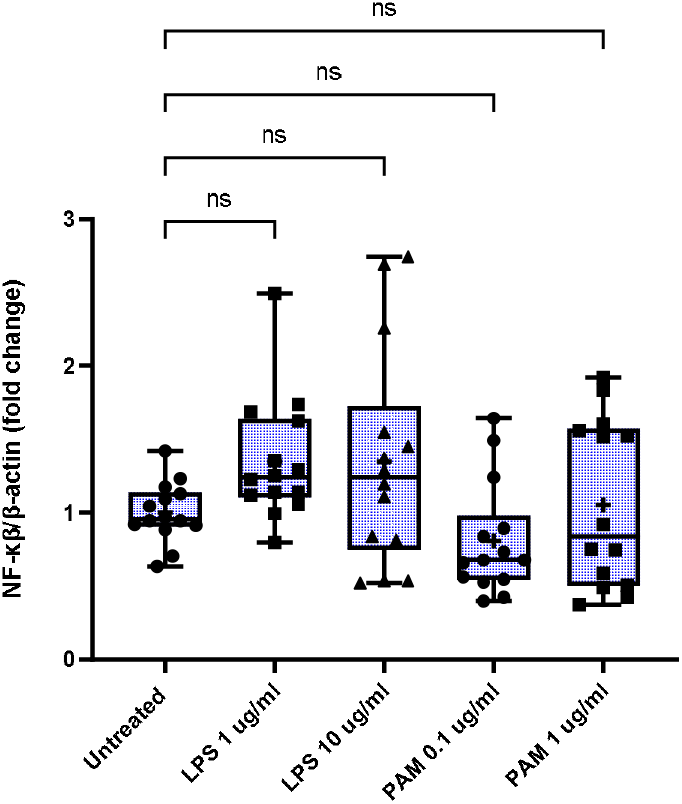
Stimulation of STC-1 cells with LPS or PAM at high or low concentrations for 24 hours had no significant effect on NF-κβ mRNA. Data are presented as box (25^th^ and 75^th^ percentiles) and whisker (5^th^ and 95^th^ percentiles), median (line), mean (+) and all data points.

**Supplementary figure 4.**
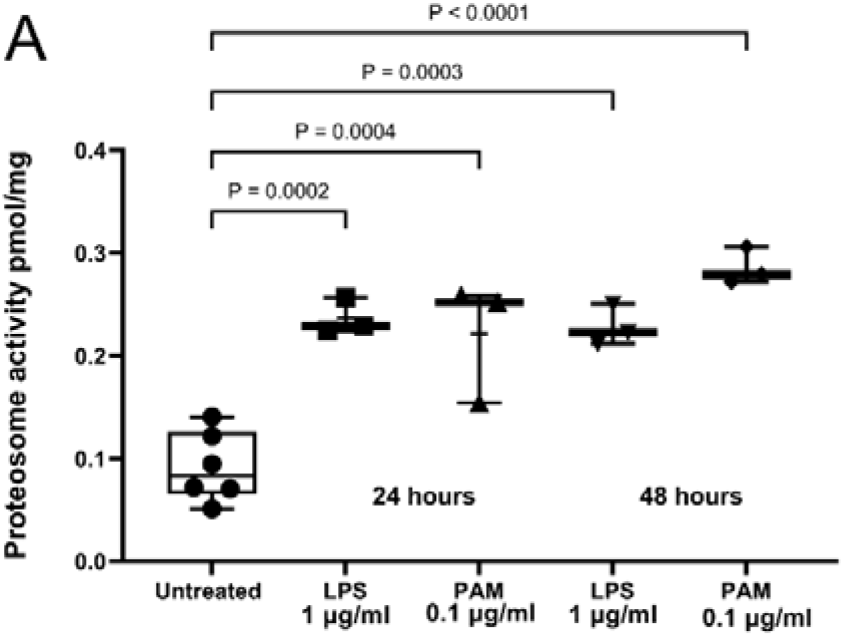
TLR agonists activate the ubiquitin-proteosome system TLR agonists increased proteosome activity (**A**) Stimulation with LPS 1 ug/ml significantly increased proteosome activity after 24 hours (P = 0.0002) and 48 hours (P = 0.0004). Stimulation with 0.1 ug/ml PAM significantly increased proteosome activity after 24 hours (P = 0.0003) and 48 hours P < 0.0001). Data are presented as box (25^th^ and 75^th^ percentiles) and whisker (5^th^ and 95^th^ percentiles), median (line), mean (+) with all data points.

